# Functional insights of two MATE transporters from *Vibrio fluvialis*

**DOI:** 10.1101/2021.01.02.425065

**Authors:** Priyabrata Mohanty, Aneri Shah, Ashima Kushwaha Bhardwaj

## Abstract

Functional characterization of H- and D-MATE (Multidrug and Toxin Extrusion) transporters from clinical isolates of *Vibrio fluvialis* revealed H-type conferred resistance to fluoroquinolones, ethidium bromide and safranin whereas D-type exhibited marginal resistance towards ethidium bromide only. Both H-/D-type transporters were inhibited by reserpine resulting in increased intracellular norfloxacin concentration. The efflux was facilitated by both Na^+^/K^+^ ions, suggesting that these efflux pumps were ion-dependent. In presence of various classes of EPIs, there was decrease in MIC exhibited by H-/D-type efflux pumps towards norfloxacin which didn’t translate into transport inhibition. But reserpine presented a conclusive pattern with decrease in MIC towards norfloxacin and increased norfloxacin accumulation inferring maximum inhibition. Substrate binding and electrostatic charge distribution of both the transporters was similar to other known MATE transporters. The H-type exhibited 10 transmembranes and D-type exhibited 11 TMs which was different from other MATE transporters known to have 12 TMs (Transmembranes). Data derived from molecular docking and ion binding studies revealed that Aspartic Acid residue in 1^st^ TM acts as ion binding site with transport mechanism similar to NorM. Electrostatic potential map of both the transporters revealed that there is a cavity formation within the transporters surrounded by charged electronegative amino acid residues. Interestingly, surface models of both transporters revealed that 1^st^ TM forms covalent bond with 7^th^ TM towards extracellular space. Docking studies also revealed that reserpine covalently binds to central pocket of both transporters and serves as excellent EPI against these transporters as evidenced by MIC and drug accumulation assays.

## Introduction

Resistance mediated by efflux pumps is one of the non-specific mechanisms that pathogenic microbes employ to lower intracellular drug concentration (Bhardwaj & Mohanty, 2012, Van Bambeke *et al.*, 2003, Poole, 2000, Poole, 2004, Poole, 2005). Efflux pumps could be primary transporters utilizing ATP for transport or could be secondary transporters that utilize sodium-/proton-motive force for transport of substrates. Till date six families of transporters are known for their involvement in multidrug resistance (MDR) in prokaryotes (Bhardwaj & Mohanty, 2012, Van Bambeke *et al.*, 2003, Hassan *et al.*, 2015). Multidrug and toxic compound extrusion (MATE) family of efflux pumps is one of the newest and least characterized family of transporters that are proven to be involved in resistance to various antibiotics and dyes including quinolones (Mohanty *et al.*, 2012, Omote *et al.*, 2006). NorM, the prototype member was discovered from *Vibrio parahaemolyticus* (Morita *et al.*, 1998) and subsequently its homologs and orthologs were identified in *Escherichia coli*, *Neisseria gonorrhoeae* and *V. cholerae* (Long *et al.*, 2008, Morita *et al.*, 2000). The MATE family further consists of three protein clusters: the NorM branch involved in MDR, a branch containing several eukaryotic proteins and a branch containing DinF involved in SOS response and MDR (Otsuka *et al.*, 2005). Growing academic interest in this family of transporters is evident from the fact that structures have been solved for a few within a span of last few years from prokaryotic as well as eukaryotic sources(Tanaka *et al.*, 2013, He *et al.*, 2010, Lu *et al.*, 2013b, Lu *et al.*, 2013a, Wong *et al.*, 2014, Tanaka *et al.*, 2017, Miyauchi *et al.*, 2017). Preliminary *in-silico* and drug transport studies of MATE family of efflux pumps revealed that these efflux pumps are sodium-ion coupled and recognize wide spectrum of substrates but neither of these characteristics have been explicitly explained at molecular level (He *et al.*, 2004). MATE pumps originated from flippases (Meeske *et al.*, 2015). Both MATE and flippases belong to same family of proteins multidrug/oligosaccharidyl-lipid/polysaccharide (MOP) transporter superfamily. Divergence can be seen as functional role has been attributed to aspartic acid in 1^st^ transmembrane of MATE-pumps, which acts as ion binding site whereas serine at 17 position is known to play functional role in MurJ flippases (Kuk *et al.*, 2017). From our laboratory, two MATE-type members (H- and D-) were first identified in *Vibrio fluvialis* (causes cholera-like illness) (Mohanty *et al.*, 2012). The genes were cloned in pBR322 vector and the protein functions were assessed using MIC and drug transport assays. Both the pumps conferred resistance to fluoroquinolones norfloxacin and ciprofloxacin. In the present study, we cloned both the transporters in arabinose promoter-based pBAD vector which allowed visualization of recombinant protein and better control over protein expression. Functional characterization was performed with assays involving MICs, drug uptake and efflux pump inhibitors (EPIs). Molecular docking, ion binding site predictions and surface electrostatics of these transporters using the known MATE-type transporters as reference proteins was carried out to further characterize the functionality of these transporters.

## Results and Discussion

### Expression of recombinant H- and D- MATE-type efflux pumps in E. coli

The sequenced genes of D- and H-type efflux pumps were designated as *vcd*, *vfd*, *vch* and vfh, *where* c stands for *V. cholerae* and *f* stands for *V. fluvialis*. Corresponding to each gene, the proteins were designated as VCH, VCD, VFH and VFD. Further the genes were cloned in arabinose driven pBAD expression system and induced as mentioned in materials in methods. Coomassie blue stained gel of arabinose-induced pBAD-*vfd* samples did not show the band corresponding to recombinant VFD at the expected position of 55 kDa (Fig. 1A) whereas the expression of positive control Lac-His was detected at 125 kDa as expected.

**Fig. 1.**
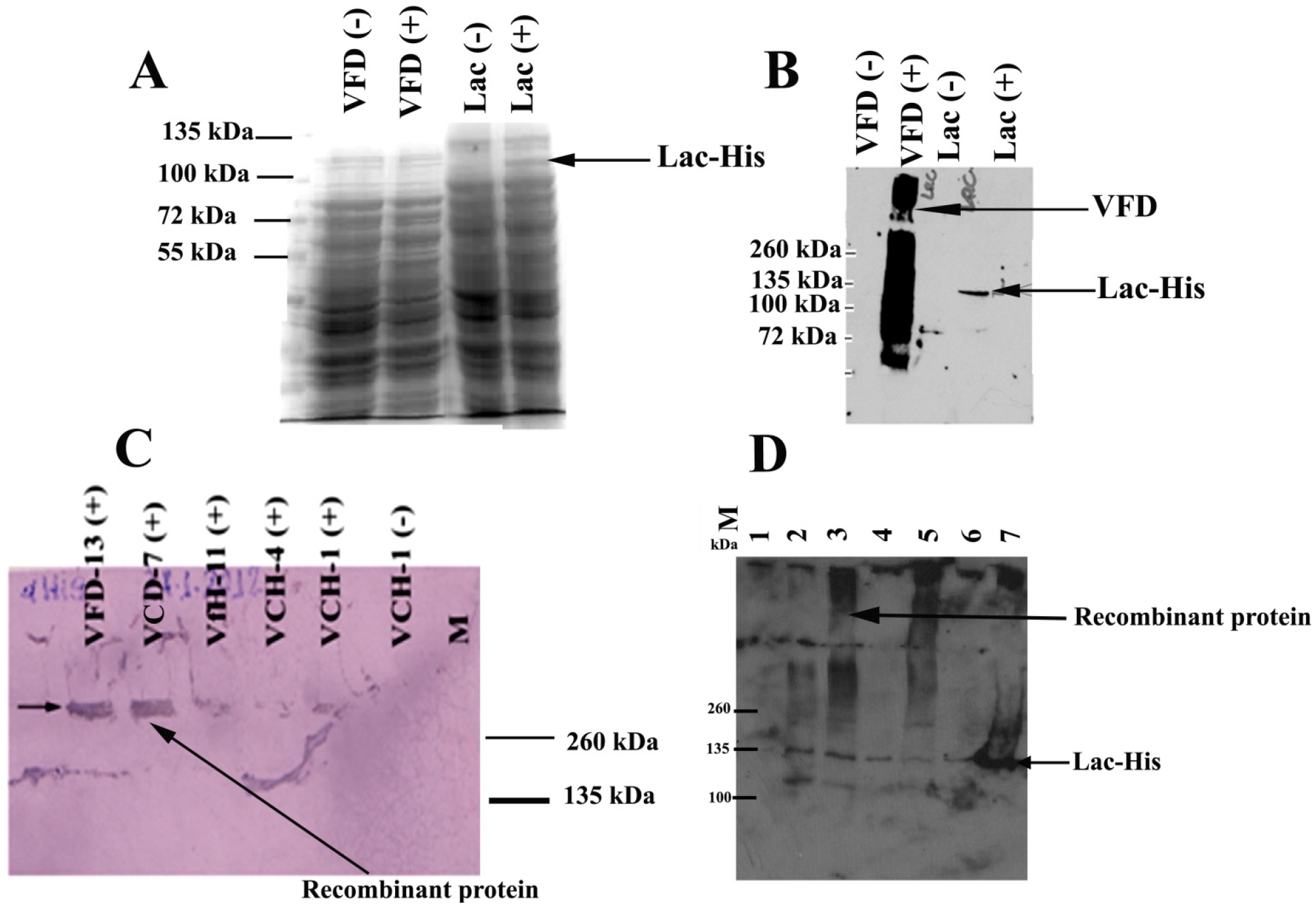
Total cell expression in induced (+) and uninduced (−) cell lysates of different recombinant pBAD clones. **A**. Coomassie blue stained 10% SDS-PAGE for total cell lysates from pBAD-*vfd* along with pBAD-Lac-His positive control in *E. coli* LMG194; **B.** Western blot of induced (+) and uninduced (−) cell lysates of pBAD-*vfd* along with pBAD-Lac-His positive control. **C.** Western blot of recombinant VCH, VFH, VCD, VFD and Lac-His in LMG194. **D.** Western blot of recombinant proteins in K-12 Δ*tolC*. The cells were induced and developed as described in materials and methods. Lane-1 Prestained Spectramulticolour (Fermentas) molecular weight markers have been indicated on the left of each pane. Lane 2,3,4 and 5 represents VCH, VFH, VCD and VFDFor Panels **B** and **D, the** blots were developed with ECL substrates (Novex, Invitrogen) and for Panel **C**, they were developed with DAB and H_2_O_2_ Goat Ant-V_5_ antibody was used as primary antibody.

Therefore, Western Blot was carried out to determine the expression of recombinant VFD protein with goat anti-V5 primary antibody and developed with enhanced chemiluminescence substrates. Results revealed the expression of Lac-His positive control at right molecular weight but interesting observation was made in case of pBAD-*vfd*, which showed diffused pattern of bands in the whole lane (Fig 4.16 B). Subsequently, all the four recombinant clones for H- and D-genes were expressed in *E. coli* LMG194 under same conditions. The development of Western blots with goat anti-His antibody using colorimetry clearly depicted a single major band with the mobility greater than 260 kDa in the lanes with induced cells for all the clones (Fig 1 C). Therefore, the slow development of the Western blots showed single major bands whereas the fast/more sensitive ECL detection could also depict the degradation pattern for the recombinant proteins.

Further, all the four recombinant clones for H- and D-genes (*vch*, *vfh*, *vcd* and *vfd*) were expressed in *E. coli* LMG194 and Western blots using colorimetry clearly depicted a single major band with the mobility greater than 260 kDa instead of 55 kDa in the lanes with induced cells for all the clones (Fig. 1C). The sensitivity of ECL led to the visualization of diffused band pattern probably indicating either various stages of oligomerisation/recombinant protein degradation whereas the less sensitive colorimetry showed a single band on Western blot. The diffused band pattern of the recombinant efflux pumps observed in the current study could be the final outcome of aberrant mobility due to oligomerisation. The observed aberrant mobility of recombinant transporter proteins due to oligomerisation has also been reported in case of ammonium transporter AmtB of *E. coli* (Blakey *et al.*, 2002) where the 45 kDa protein showed series of bands of varying molecular weights suggesting it to be an oligomer. In our study, the mobility of the H- and D-recombinant effux pump proteins showed similar pattern on Western blots where ~55 kDa proteins migrated at >260 kDa molecular weight suggesting that the recombinant protein existed as oligomers that were stable in the presence of SDS and β-ME. The recombinant efflux pumps formed a series of aggregated species due to the presence of hydrophobhic patches that appeared as a ladder of high molecular weight bands under denaturing and reducing conditions (Blakey *et al.*, 2002). The wild type Na^+^/H^+^ antiporter of *Methanococcus jannaschii* presents itself as a dimer even in the presence of SDS and undergoes proteolytic degradation (Goswami *et al.*, 2011). The recombinant expression of membrane proteins is tricky where specialized expression systems have been developed to counteract protein degradation and oligomerisation (Wagner *et al.*, 2008). Taken together, the facts from earlier studies and observations from the present study suggested that H- and D-MATE efflux pumps exhibited aberrant mobility possibly be due to oligomerisation similar to AcrB due to intra-molecular van der Waals bonding that resists dissociation in presence of SDS and β-ME.

### pBAD recombinant efflux pumps are translocated to the membrane

Upon successful detection of recombinant proteins in the host cells with aberrant mobility, classical traits of membrane proteins, further studies were carried out to determine if the recombinant proteins were actually targeted to the membrane of the *E. coli* host. Two approaches *i.e.* confocal microscopy and membrane fractionation were utilised to investigate the same.

Earlier bioinformatics studies of both the efflux pumps had predicted that these pumps were transmembrane proteins (Mohanty *et al.*, 2012). Localization of these efflux pumps in cell membrane was investigated utilizing confocal microscopy where the laser was programmed to detect membrane marker Film Tracer 1-43 and CY3 fluorochrome conjugated secondary antibody from the same pin hole. Results revealed that VFD and VFH were localised in the membrane of *E. coli* LMG194 (Fig. 2 Panel A: A-F). The recombinant proteins generated red signal due to CY3 (Fig. 2 A, panelsB and E) and membrane marker generated green signal for cell membrane (Fig. 2A,panels A and D). DAPI, which was used for nuclear staining was also employed as counterstain but it did not bind with the nucleoid and was not visible by confocal imaging. Change in the colour of membrane to yellow in the superimposed image of recombinant efflux pump protein and membrane marker (Fig. 2A, panels C and F) suggested that the recombinant proteins were localised in the membrane and were detected by the same pinhole of the microscope.

**Fig. 2.**
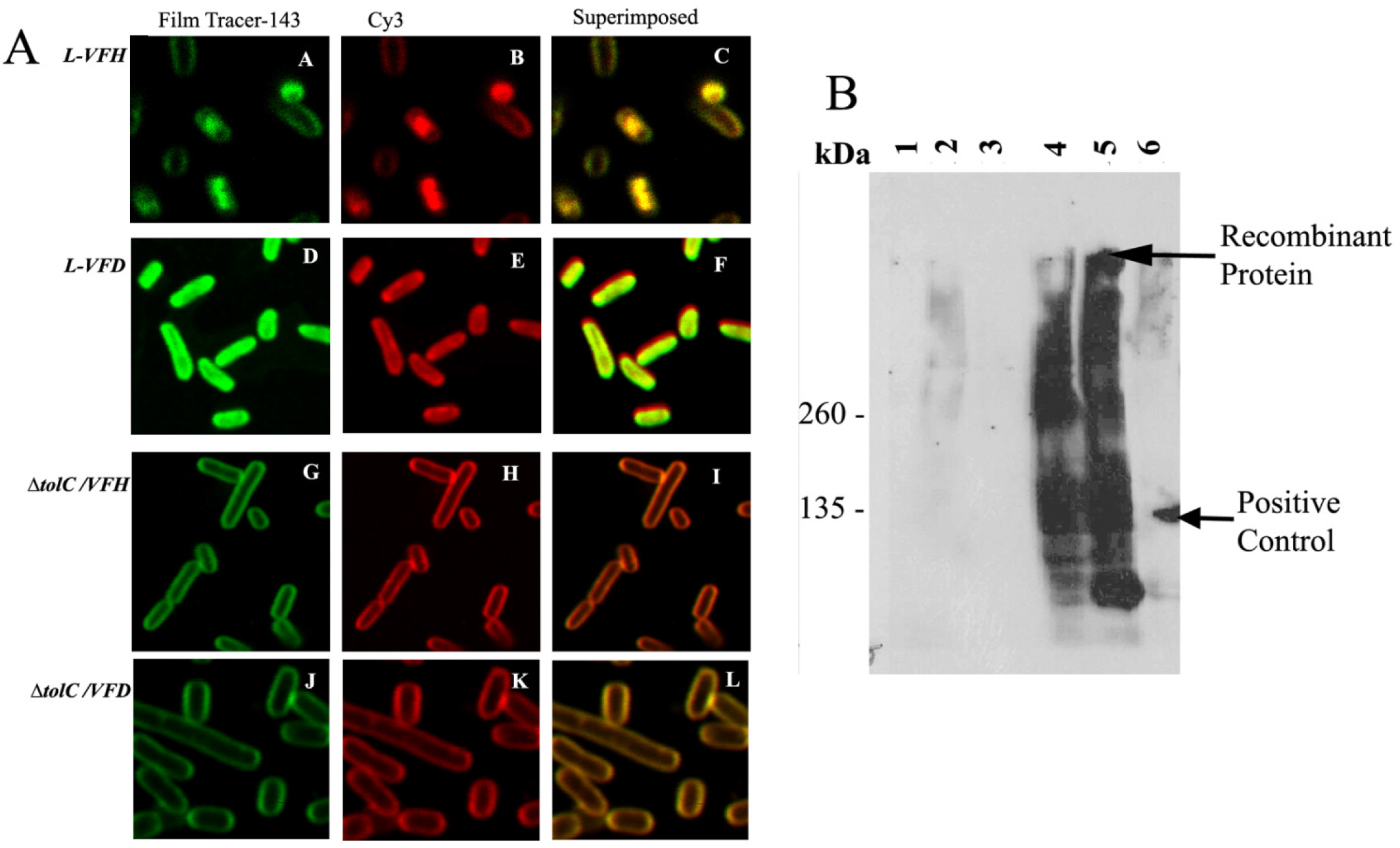
Membrane localisation of recombinant efflux pumps in *E. coli.* **Panel A.** Confocal imaging of pBAD-recombinants in heterologous *E. coli* host. **A/D**:Visualisation of *E. coli* cells harbouring pBAD-*vfh/vfd* stained with membrane marker Film-tracer 1-43; **B/E**:Detection of recombinant pBAD-*vfh/vfd* in *E. coli* labelled with Goat anti-Mouse Cy3 secondary antibody; **C/F** (63X ZOOM 4): Superimposed/merged images from **A/B or D/E** of recombinant pBAD-*vfh/vfd* in *E. coli* LMG194; **G/J**: Visualisation of *E. coli* cells harbouring pBAD-*vfh/vfd* using membrane marker Film-tracer 1-43; **H/K**: Detection of recombinant pBAD-*vfh/vfd* in *E. coli* labelled with Goat anti-Mouse Cy3 secondary antibody; **I/L**: Superimposed/merged images from **G/H** or **J/K** of recombinant pBAD-*vfh/vfd* in *E. coli* K-12 Δ*tolC*2 Δ*tolC*. **Panel B.** Membrane fractionation of *E. coli* harbouring pBAD-*vfd.* 63X Zoom 4 settings were used for obtaining these images. M- Spectramulticolour protein marker (Fermentas). Lane 1- VFD −ve- Total cell lysates for uninduced samples; Lane 2-VFD +ve- Total cell lysates for induced samples; Lane 3- VFD Son (Sup) - total soluble fraction after sonication.; Lane 4- VFD SAR (Sup)- Total soluble proteins after sarkosyl treatment; and Lane 5- VFD SAR Pellet- debris and other subcellular organelles. Lane 6- Lac-HisPositive control. Goat anti-V5 antibody was used as primary antibody. HRP conjugated Mouse Anti-Goat IgG was used as secondary antibody (Jackson Immunoresearch). Prestained Spectramulticolour (Fermentas) molecular weight marker in kDa has been indicated of the left of panel. The blots were developed with Novex ECL substrates (Invitrogen).

To further confirm if the recombinant proteins were localized in the inner membrane, a protocol utilising a combination of membrane fractionation and selective solubilisation of the inner membrane with detergents was employed (Hobb *et al.*, 2009, Leyh, 1990, Frankel *et al.*, 1991).Western blot revealed that upon induction, the recombinant protein was detected in total cell lysate (Fig. 2B, lanes 1 and 2) for VFD. The fraction containing total soluble protein did not show the presence of recombinant protein indicating that the recombinant protein was not cytosolic but probably associated with insoluble fraction (Fig. 2B, lane 3). Sarkosyl solubilised inner membrane fraction (lane 4) revealed the band pattern corresponding to recombinant VFD confirming that the protein was translocated to the inner membrane. This protein was also detected in the sarkosyl treated outer membrane pellet indicating partial solubilisation of the membrane upon sarkosyl treatment (Fig. 2B, lane 5). Apart from the above, partially purified recombinant VFD from inner membrane also showed aberrant migration and diffused band pattern as observed earlier.

### MIC assays for pBAD recombinants in E. coli K-12 ΔtolC indicated different substrate profile for H and D pumps

The recombinant constructs for H- and D-type proteins when introduced into *E. coli* LMG194 did not exhibit any change in the MIC against ethidium bromide and norfloxacin although both the compounds had been shown as substrates for these membrane proteins in earlier studies from this laboratory (Mohanty *et al.*, 2012). Earlier studies with pBR322 expression system had utilized *E. coli* host KAM32 with deletion of two major efflux pumps AcrAB and YdhE for functional characterization of the recombinant proteins expressed in this host (Mohanty *et al.*, 2012). In the current study *E. coli* K-12 strains with deletions in genes for major RND efflux systems; Δ*acrA* (deletion of connecting protein), Δ*acrB* (deletion of its inner protein) and Δ*tolC* (deletion of its outer membrane protein) were utilized for functional characterization of pBAD recombinants (Kuroda & Tsuchiya, 2009, Matsuo *et al.*, 2008, Matsuo *et al.*, 2007, Otsuka *et al.*, 2005, Mohanty *et al.*, 2012).

Results for MICs of ethidium bromide, ciprofloxacin and norfloxacin showed that though all the three K-12 deletion strains, K-12 Δ*acrA*, K-12 Δ*acrB* and K-12 Δ*tolC* showed reduction in MIC as compared to the control wild type strains K-12 and LMG194, K-12 Δ*tolC* exhibited maximum hypersusceptibilty *i.e.* at least 16-fold decrease in MIC towards ethidium bromide and norfloxacin and more than 8-fold decrease in MIC towards ciprofloxacin (Supplementary Table 1). Subsequently, accumulation assays also revealed that *E. coli* K-12Δ*tolC* had the highest drug accumulation in comparison to the other hosts utilised in the study (Supplementary Fig.1).

Western blot and confocal microscopy (Fig. 1D and Fig. 2A, panels G to L respectively) revealed that the recombinant proteins were expressed in K-12Δ*tolC* and localized in its inner membrane. Taken together, the results of Western blots, confocal microscopy, MIC and accumulation assays (Supplementary Fig.1) indicated that K-12 Δ*tolC* was a suitable hypersusceptible strain for expression and functional characterization of the recombinant MATE pumps.

Subsequently, *E. coli* K-12 Δ*tolC* expressing pBAD recombinants for VFH, VCH, VFD and VCD presented an interesting profile in terms of recognition/transport by these two pumps (Table 1). H-type pumps exhibited increase in MIC for ethidium bromide (at least 4-fold), norfloxacin and ciprofloxacin (atleast 2-4-fold), acriflavin (2-fold) and safranin (4-fold). As against this, D-type pumps conferred resistance only towards ethidium bromide with at least 2-fold increase in MIC when compared to the control. Both H- and D-type pumps showed almost similar/slight decrease (2-fold) in MIC for the aminoglycosides (neomycin and gentamicin). Both H- and D-type pumps showed unaltered MIC for nalidixic acid, ofloxacin and acridine orange while D-type pumps additionally showed unaltered MIC for norfloxacin, ciprofloxacin, acriflavin and safranin.

**Table 1.** MIC for recombinant clones carrying MATE-type efflux pump genes from *V. fluvialis* and *V. cholerae* in pBAD expression vector and *E. coli* Δ*tolC* host.

| | MIC in $\mu\text{g/mL}$ for the recombinant clones | | | | |
| --- | --- | --- | --- | --- | --- |
| Drugs/ Dyes | $\Delta tolC$ | $\Delta tolC$ /VCH | $\Delta tolC$ /VFH | $\Delta tolC$ /VCD | $\Delta tolC$ /VFD |
| Ethidium Bromide | <3.9 | **15.6** | **15.6** | **3.9** | **7.8** |
| Norfloxacin | <0.0156 | **0.03125-0.0625** | **0.03125-0.0625** | <0.0156 | <0.0156 |
| Ciprofloxacin | <0.00156 | **0.003125-0.00625** | **0.003125-0.00625** | <0.00156 | <0.00156 |
| Nalidixic Acid | 0.3125-0.625 | 0.3125-0.625 | 0.3125-0.625 | 0.3125-0.625 | 0.3125-0.625 |
| Ofloxacin | 0.0125-0.025 | 0.0125-0.025 | 0.0125-0.025 | 0.0125-0.025 | 0.0125-0.025 |
| Neomycin | >200 | 200 | 200 | 200 | 200 |
| Gentamicin | 3.125 | 1.56-3.125 | 3.125 | 1.56 | 1.56 |
| Acridine Orange | 37.5 | 37.5 | 37.5 | 37.5 | 37.5 |
| Acriflavin | 6.25 | **12.5** | **12.5** | 6.25 | 6.25 |
| Safranin | 3.125 | **12.5** | **12.5** | 3.125 | 3.125 |

As kanamycin cassette is inserted in place of the gene of interest ( $\Delta tolC$ ) to be disrupted, so MIC against kanamycin could not be evaluated against pBAD recombinants in $\Delta tolC$ . Increase in MIC is shown as bold.

NorM, the prototype member of MATE-type efflux pumps from *V. parahaemolyticus* is a broad spectrum efflux pump reported to confer resistance towards basic lipophilic compounds like ethidium bromide and rhodamine, hydrophilic aminoglycosides like kanamycin and neomycin and hydrophilic zwitterionic fluoroquinolones such as norfloxacin and ciprofloxacin, but not to hydrophobic quinolones such as sparfloxacin and nalidixic acid (Morita *et al.*, 1998). In the present study, H-type recombinant seemed to be an efflux pump with primary transport capability towards zwitterionic fluoroquinolones and basic lipophilic compounds. Earlier studies had revealed that H-type pumps (VCH/VFH) exhibited resistance towards norfloxacin and ciprofloxacin only when expressed in pBR322 expression system (Mohanty *et al.*, 2012) but when these pumps were expressed in pBAD/Δ*tolC* system, they exhibited norfloxacin and ciprofloxacin resistance along with extended resistance towards ethidium bromide, acriflavin and safranin. The expression of efflux pumps is generally very low and results in a modest increase of two-four folds in MIC (Long *et al.*, 2008) which was also observed in the current study (Long *et al.*, 2008). KetM from *Klebsiella pneumoniae* also exhibited narrow spectrum resistance to fluoroquinolones with modest change in MIC (Ogawa *et al.*, 2015). Asp32, Glu251 and Asp367 from *V. parahaemolyticus* NorM were implicated in sodium-dependent transport of antimicrobials and Asp32 specifically is known for its role in efflux thus conferring resistance towards norfloxacin and ethidium bromide (Otsuka *et al.*, 2005). Even though the efflux pumps in the present study carried only Asp at position 40/38 in TM1 out of three conserved residues they were able to exhibit marginal MDR phenotype. This also gives a notion that apart from these three amino acids, other residues might play important role in transport. It was later proven in MATE transporter from the plant *Camelina sativa* (Tanaka *et al.*, 2017). Glutamic Acid, glutamine, asparagine, arginine along with aspartic acid amino acid residues were implicated for their role in forming hydrophobic cavity formation and transport in MATE transporter from the plant *Camelina sativa* (Tanaka *et al.*, 2017).

### Accumulation assays revealed that the recombinant strains were able to efflux drugs/dyes

The substrate transport studies revealed that there was an increased accumulation of ethidium bromide by pBAD recombinants expressing either H-/D-type efflux pumps in comparison with control as also observed in earlier studies with their corresponding pBR322 recombinants (Mohanty *et al.*, 2012) (Fig. 3A and 3B).

**Fig. 3.**
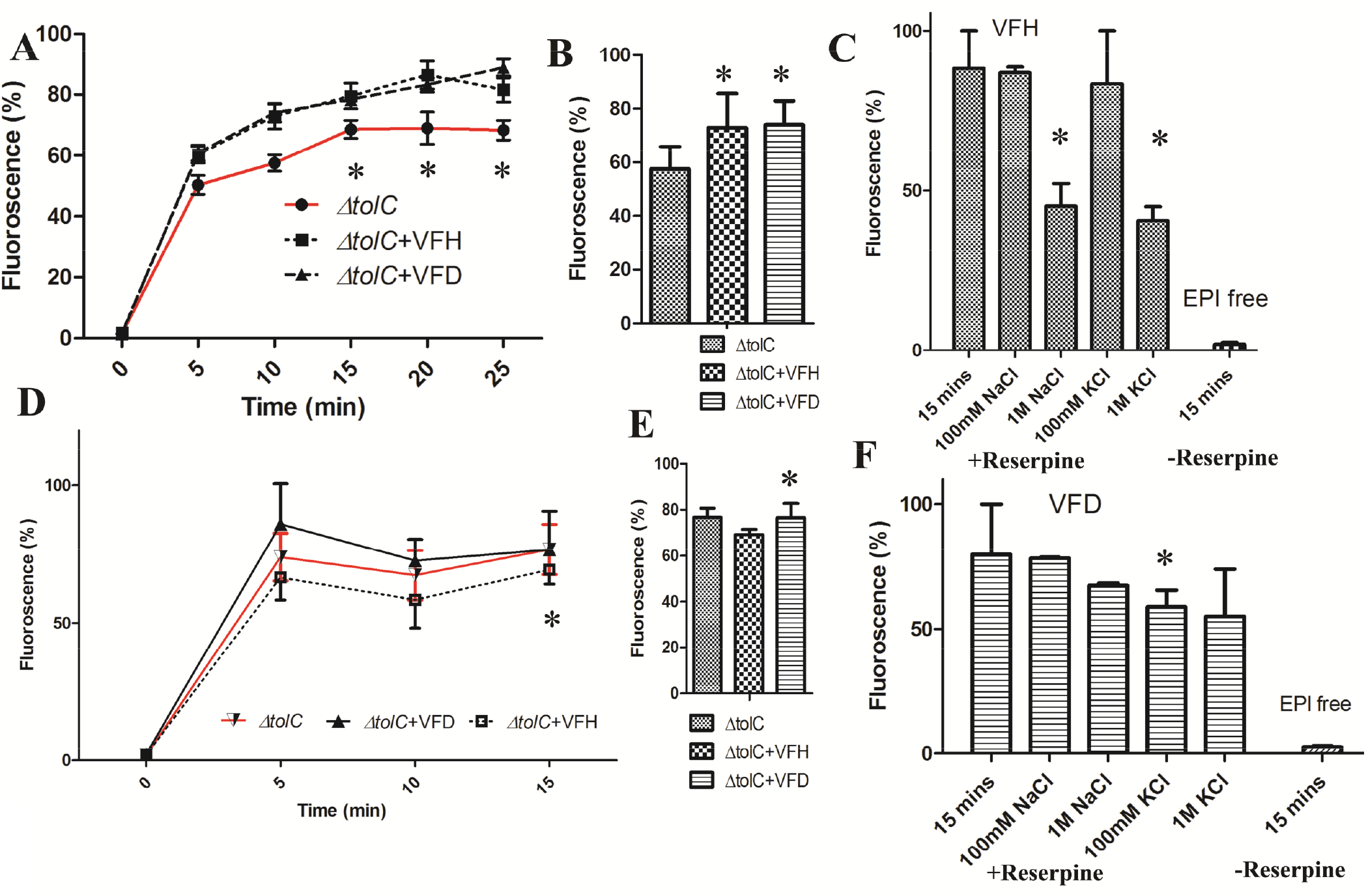
**A/D.** Intracellular ethidium bromide/norfloxacin accumulation in recombinant K-12 Δ*tolC* harbouring *vfd* and *vfh* genes in pBAD vector. Samples were collected at every 5 mins for processing and fluorescence was measured A/**B.** Intracellular ethidium bromide accumulation in recombinant K-12 ΔtolC harbouring *vfd* and *vfh* genes in pBAD vector after 15 mins. D/**E.** Intracellular norfloxacin accumulation in recombinant K-12 ΔtolC harbouring *vfd* and *vfh* genes in pBAD vector after 15 mins. **C/F.** Ion dependence of pBAD recombinant efflux pumps in the presence of 100 mM or 1 M sodium chloride (NaCl) or 100 mM or 1 M potassium chloride (KCl) in K-12 ΔtolC cells transformed with *pBAD-vfh/vfdVFD*. Data is expressed as mean percentage fluorescence intensity of the maximum fluorescence intensity obtained during the experiment from three independent experiments and * represents significant difference to control (*P* < 0.05).

**Fig. 4.**
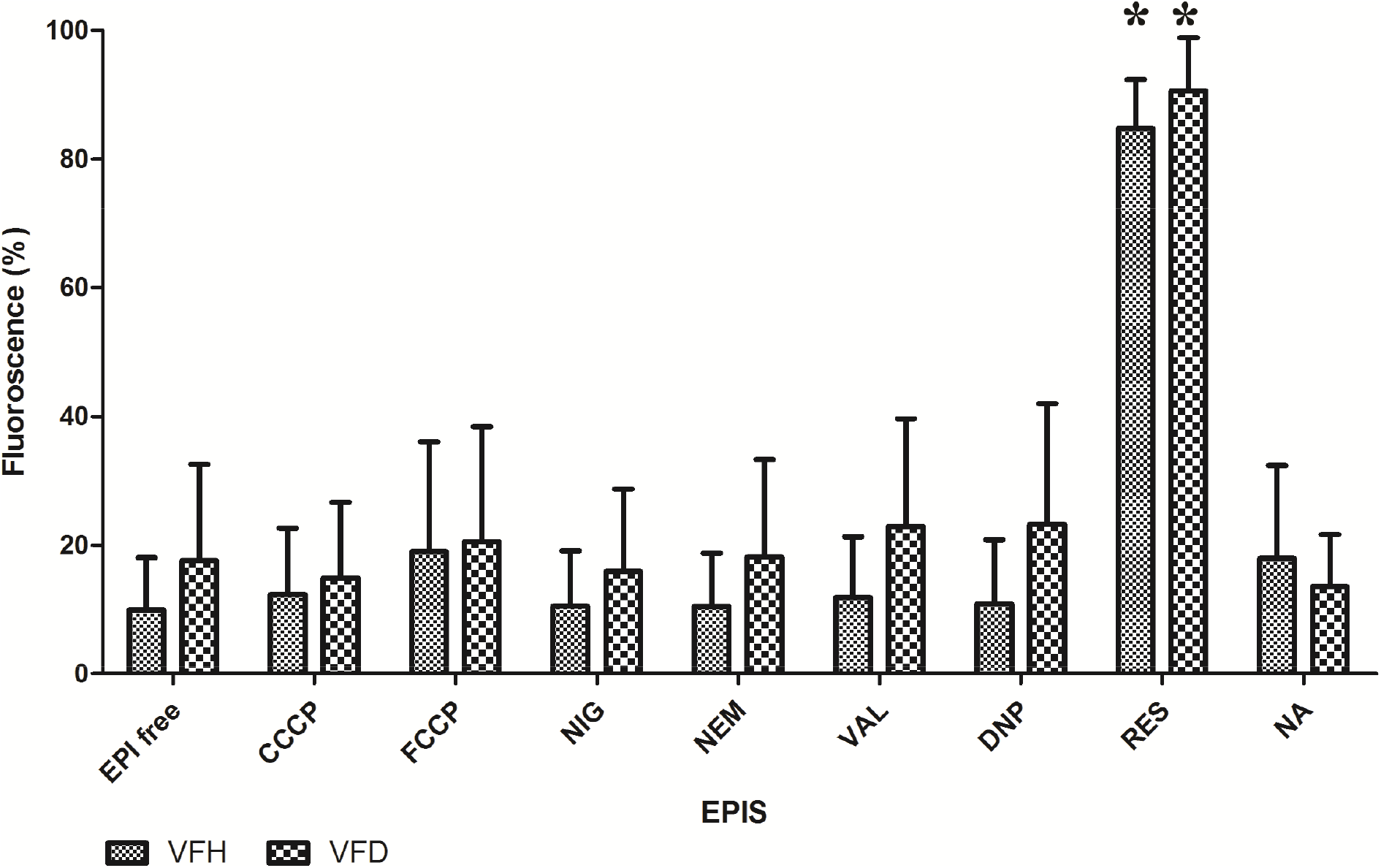
Accumulation of norfloxacin in presence of various EPIs. Accumulation of norfloxacin by *E. coli* K-12 Δ*tolC* expressing H- and D-type pumps in the presence of EPIs. Recombinant *E. coli* K-12 ΔtolC was loaded with norfloxacin and incubated at 37°C for 15 mins in presence/absence of EPIs. These samples were subsequently processed and analyzed for fluorescence intensity as described in the materials and methods. * represents significant difference to control (*P* < 0.05). The concentration of EPIs used in μg/mL are FCCP 3.125, CCCP 1.56, NIG 50, VAL 50, NEM 62.5, DNP 25, RES 125 and NA 50.

The intracellular ethidium bromide accumulation profiles of both the efflux pumps was in contrast to the results of MIC assays where elevation in MIC was observed towards ethidium bromide (Fig. 3A and 3B; Table 2). The absence of correlation between the results of MIC and accumulation assays particularly in ethidium bromide could result due to the time period of both the assays.

**Table 2.**
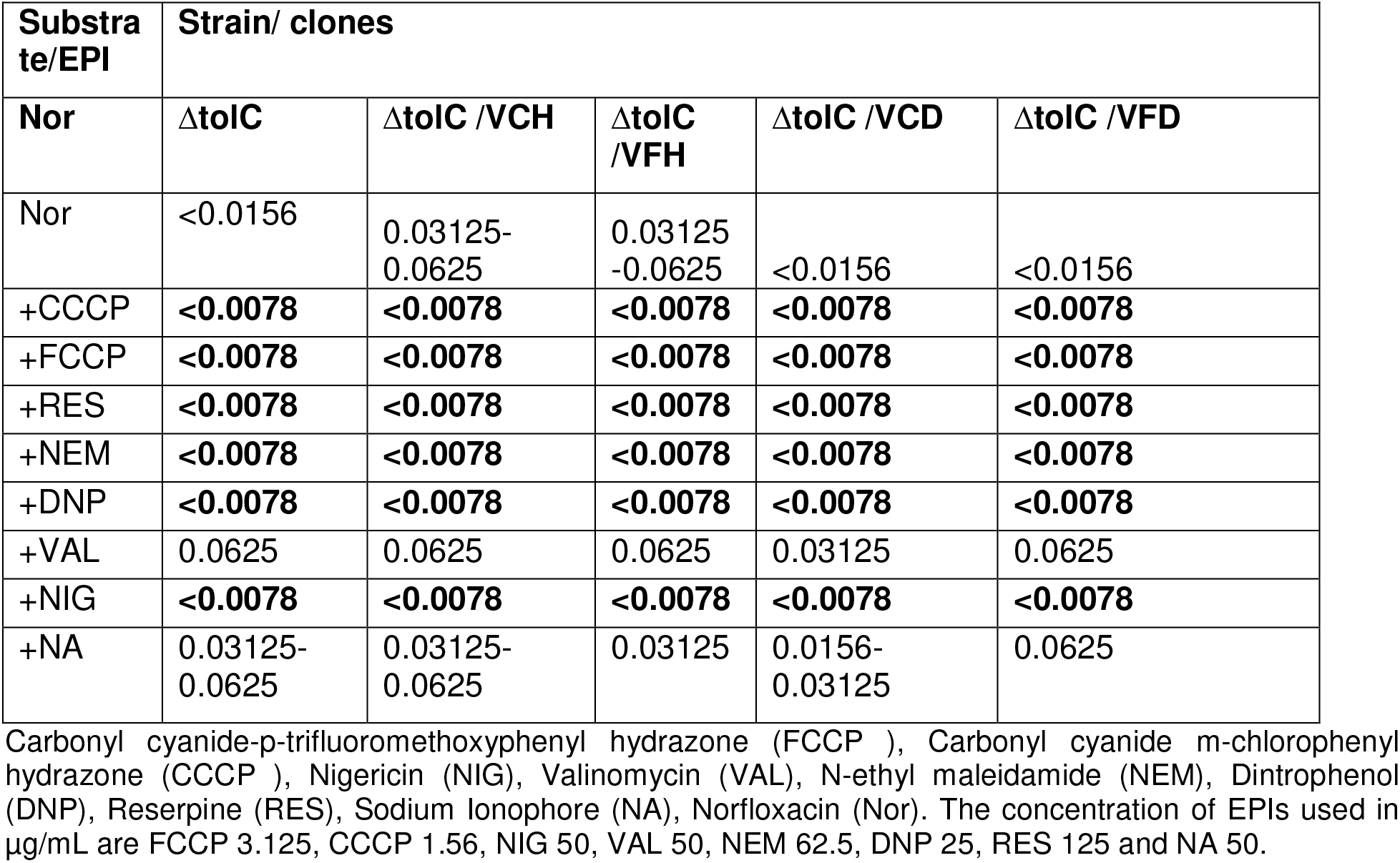
Change in MICs of norfloxacin in pBAD recombinant clones carrying MATE-type efflux pump genes from *V. fluvialis* and *V. cholerae* with addition of EPIs.

However, in case of norfloxacin, there was a correlation between MIC assays and accumulation assays. Results revealed that recombinants with H-type pumps which exhibited increased MIC towards norfloxacin, showed lowering of intracellular norfloxacin accumulation whereas recombinants harbouring D-type efflux pumps that exhibited unaltered susceptibility towards norfloxacin showed almost similar intracellular norfloxacin concentration as compared to the control (Fig. 3D and 3E). Subsequently ion dependence assays were performed for both the transporters to understand their fueling mechanism.

### The Na^+^/K^+^-ions equally facilitated the efflux of norfloxacin by H- pumps

Earlier studies from our laboratory as well as others had revealed that MATE pumps belonging to NorM cluster could transport their substrates in the presence of monovalent ions like Na^+^ (Morita *et al.*, 1998, Mohanty *et al.*, 2012). The K-12 Δ*tolC/* pBAD-*vfh* recombinants were loaded with norfloxacin in the presence of reserpine, a compound known to block efflux activity. Subsequently, different concentrations of 0.1 M and 1 M NaCl or KCl were added to the assay mixture and intracellular norfloxacin was monitored to assess the effect of these ions on the efflux activity of these pumps. Results showed that there was an increase in intracellular norfloxacin accumulation in this recombinant in the presence of reserpine compared to reserpine free control (Fig. 3C and 3F, columns 1 and 6 respectively). Though the addition of 100 mM KCl/NaCl did not significantly induce the efflux, increase in salt concentration to 1M KCl/NaCl resulted in a greater efflux of the drug from these cells compared to the cells without any salts (Fig. 3C and 3F, columns 2-5).

Earlier studies on MATE- efflux pumps had established that these efflux pumps were fueled either by Na^+^/H^+^-ions for transport (Bhardwaj & Mohanty, 2012) which was also observed in the current study. There was an increase in intracellular drug concentration in H-recombinants when the Efflux Pump Inhibitor (EPI) reserpine was added. Effluxing commenced on addition of Na^+^/ K^+^ ions. Newer studies also point out that the transport capability of MATE-type *E. coli* ClbM transporter involved in transport of precolibactin was equally influenced by Na^+^, K^+^ and Rb^+^ ions (Mousa *et al.*, 2017). K^+^ ions are known to be congener of Na^+^ ions. Another study revealed that NorM from *V. parahaemolyticus* utilises both PMF and Na^+^ for transport (Jin *et al.*, 2014). Overall it seems both the transporters are Na^+^-ion dependent which has been proved earlier (Mohanty *et al.*, 2012).

### Changes in MIC of norfloxacin in the presence of EPIs

Different types of EPIs were utilised to deduce actual transport mechanism of these efflux pumps.. For this, the MIC of these EPIs was deduced against host *E. coli* K-12 Δ*tolC* and sub-lethal concentrations of EPIs (~50% of MIC) were utilized to monitor changes in MIC/drug accumulation patterns for recombinant efflux pumps.

Results revealed that compounds like FCCP and CCCP were effective at low concentrations to host and with norfloxacin a decrease in MIC was observed for H-type (at least 8-folds) and for D-type (2 folds) Table 2). Both compounds are known to kill living cells at low concentration as they disrupt PMF, and have been extensively utilised for inhibition of efflux pumps with concentrations varying for each study (Begum *et al.*, 2005, Morita *et al.*, 2000, Morita *et al.*, 1998). Other compounds like DNP which deplete ATP or compounds like NIG, NA and VAL which are known as H^+^, Na^+^ and K^+^ specific ionophores respectively, exhibited high MIC for host. But with norfloxacin, DNP and NIG also decreased the MIC by at least 2-8-folds for norfloxacin. against both the compounds (Table 2). VAL and NA did not inhibit both the pumps for norfloxacin (Table 2). These ionophores mostly increase transferability of ions across membrane involved in generating EMF, which may or may not be important for energy recycling (Castro *et al.*, 2015). RES and NEM, both of which either bind or covalently modify the efflux pumps, also exhibited very high MIC for *E. coli* K-12 ΔtolC but in conjunction with norfloxacin, both compounds decreased the MIC (Table 2). Both NEM and RES decreased the MIC exhibited by both pumps by at least 2-16-folds whereas (Table 2). It was not surprising that reserpine increased sensitivity towards fluoroquinolones. Resperine had been earlier reported to inhibit various efflux pumps resulting in increased susceptibility (Baranova & Neyfakh, 1997, Brenwald *et al.*, 1997, Aeschlimann *et al.*, 1999, Markham, 1999, Markham *et al.*, 1999, Gibbons & Udo, 2000, Mullin *et al.*, 2004).

### Accumulation of norfloxacin by H-/D-type pumps in the presence of EPIs

The MIC assays had established that some of the EPIs were active in reducing the MIC of norfloxacin for the recombinants expressing efflux pumps. Therefore, norfloxacin accumulation studies were performed to assess the real time effect on drug transport by the H-/D- recombinants in the presence/absence of EPIs.

Results revealed that *E. coli* K-12 Δ*tolC* expressing H-type pumps showed a marginal to moderate degree of increased norfloxacin accumulation in presence of CCCP and FCCP in comparison to EPI free control (Fig. 4). These pumps exhibited almost unaltered/marginal increased accumulation of norfloxacin in presence of NIG, NEM, VAL and DNP (Fig. 4). Interestingly, both efflux pumps exhibited very high accumulation of norfloxacin in presence of RES (Fig. 4). MIC studies of norfloxacin against both recombinants in the presence of EPIs had revealed that compounds like CCCP, RES and FCCP decreased the MICs (Table 4). This was in sync to the norfloxacin accumulation results, where these compounds increased the accumulation of norfloxacin. NA increased the intracellular norfloxacin for H-type of efflux which was in contrast to MIC assays where it did not decrease the MIC for norfloxacin (Table 2).

However, for NEM, DNP and NIG this correlation between MIC and accumulation results was missing probably due to the kinetics of inhibition by these EPIs and the difference in the time course for the two type of assays. MATE family of efflux pumps utilise Na^+^ ions for transport which was observed in the current study. There was an increased intracellular norfloxacin accumulation with utilisation of H^+^/ Na^+^/K^+^ ion specific ionophores such as NA, CCCP and FCCP. This was in corroboration with our earlier observation in ion dependence experiments where we observed that both the efflux pumps showed greater efflux in presence of both Na^+^ and K^+^ ions also evidenced in MIC experiments (Table 2) where these EPIs decreased the MIC of norfloxacin. Taken together, there was a relative correlation between drug accumulation, ion dependence and change in MIC patterns in case of some selected EPIs.

### Molecular Modeling and docking

Structural prediction of MATE-type efflux pumps had suggested 12 transmembrane helices in their structure which was further validated with the structure of first Na^+^ dependent MATE-type efflux pump from *Pyrococcus furiosus* having 12 transmembranes (Lomovskaya & Bostian, 2006, Tanaka *et al.*, 2013). In our earlier attempt in predicting 3D structures of VFH and VFD using I-TASSER server, we had reported that VFD had 11 transmembranes (TMs) whereas VFH had 10 TMs (Mohanty *et al.*, 2012). After 2012, more structures were solved for the members of this family of transporters (Tanaka *et al.*, 2013, He *et al.*, 2010, Lu *et al.*, 2013b, Lu *et al.*, 2013a). Subsequent to these studies, further attempts for protein modeling were made using SwissModel (https://swissmodel.expasy.org/).to understand the topology and mechanism of efflux by these two MATE pumps Both PyMol (https://pymol.org/2/) and Chimera (https://www.cgl.ucsf.edu/chimera/) were used forvisualization of their molecular structures. Results revealed that both the efflux pumps had 10 predicted TMs arranged in two bundles of 5 TMs each (Fig. 5.) similar to crystal structures of earlier published MATE transporters (Lu *et al.*, 2013b). They were also outward facing like NorM (Lu *et al.*, 2013b) with distinct internal cavity. It was both in corroboration (for VFH) and contrast (for VFD) to our earlier finding where VFH model was similar to NorM and VFD had different predicted protein model (Mohanty *et al.*, 2012). Subsequently, molecular docking was done using SwissDock (http://www.swissdock.ch/) to validate binding interactions of both the transporters with ethidium bromide and norfloxacin as both the transporters showed certain degree of transport for these substrates. Classically, a MATE pump has 12 TMs arranged in two bundles of six TMs each (Tanaka *et al.*, 2013, Tanaka *et al.*, 2017, Radchenko *et al.*, 2016, Lu *et al.*, 2013b).

**Fig. 5.**
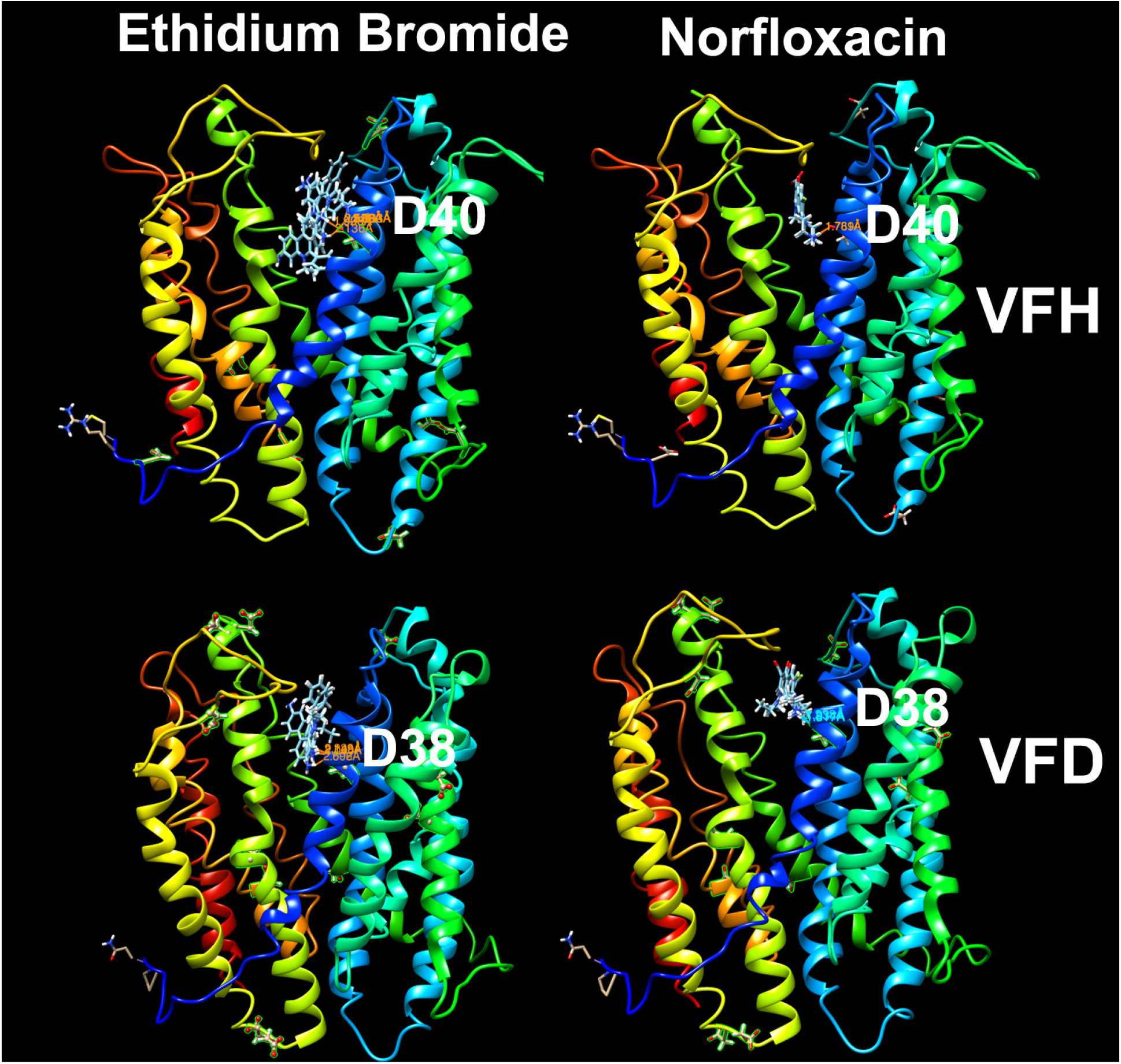
Docked pose of ethidium bromide and Norfloxacin with *V. fluvialis* MATE efflux pumps (VFH and VFD) forming covalent complex at Asp 38/40 of 1^st^ transmembrane.

The first TM has an ion binding site represented by D32, the Aspartic Acid residue at 32 position (Otsuka *et al.*, 2005, Long *et al.*, 2008, Lu *et al.*, 2013a, Lu *et al.*, 2013b). With this input, the protein sequences of both the transporters were submitted to SwissDock ≈server (http://www.swissdock.ch/). The docking was programmed for finding all possible ligand interactions. All aspartic acid residues were first identified from the generated models and hydrogen bonds were calculated. The details have been presented in Table 3 and Fig 5.

**Table 3.**
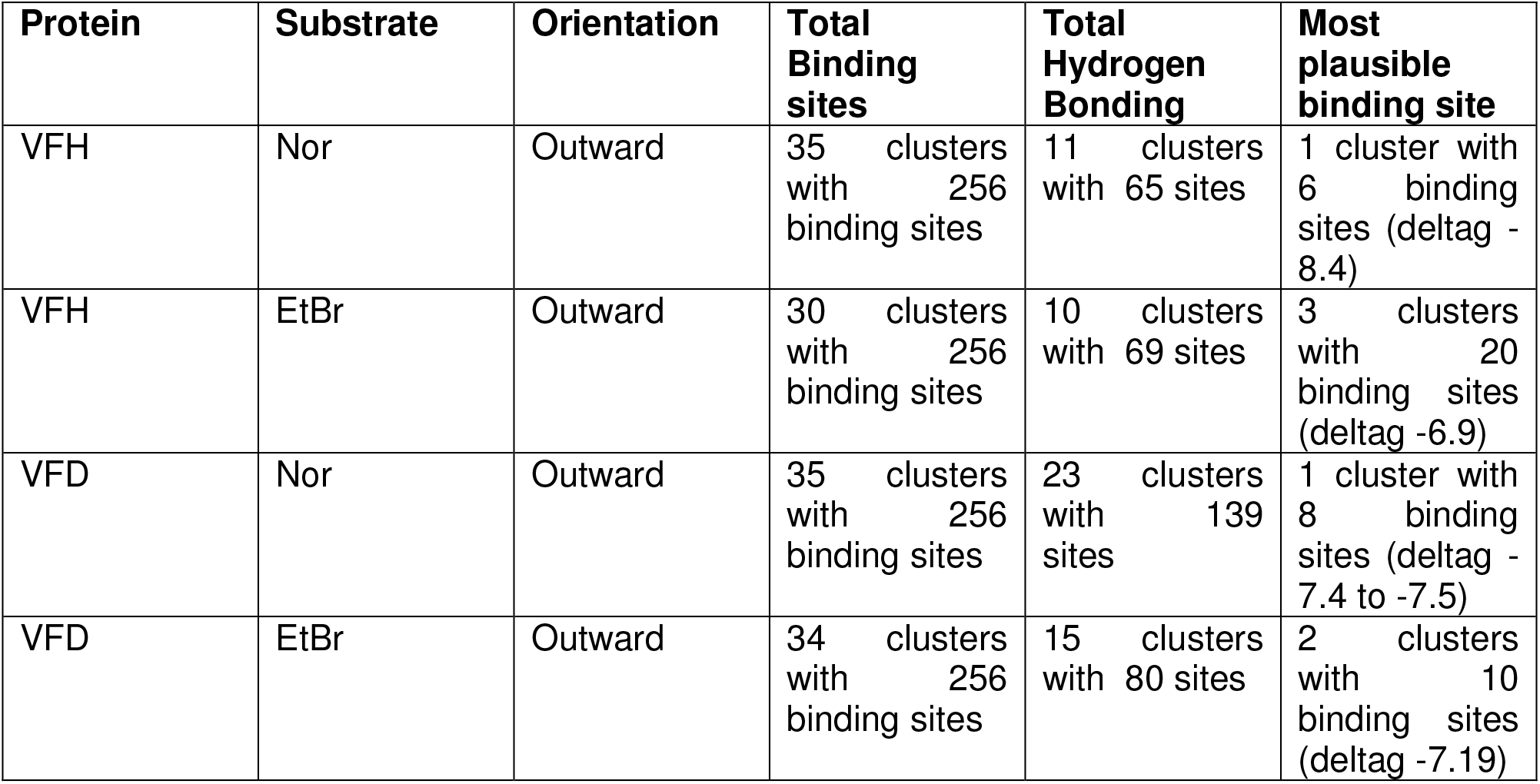
Ligand-docking of VFH and VFD efflux pumps with various substrates.

| Protein | Substrate | Orientation | Total Binding sites | Total Hydrogen Bonding | Most plausible binding site |
| --- | --- | --- | --- | --- | --- |
| VFH | Nor | Outward | 35 clusters with 256 binding sites | 11 clusters with 65 sites | 1 cluster with 6 binding sites (deltag - 8.4) |
| VFH | EtBr | Outward | 30 clusters with 256 binding sites | 10 clusters with 69 sites | 3 clusters with 20 binding sites (deltag -6.9) |
| VFD | Nor | Outward | 35 clusters with 256 binding sites | 23 clusters with 139 sites | 1 cluster with 8 binding sites (deltag - 7.4 to -7.5) |
| VFD | EtBr | Outward | 34 clusters with 256 binding sites | 15 clusters with 80 sites | 2 clusters with 10 binding sites (deltag -7.19) |

Interaction between the protein and ligands was optimum in the first TM at D40 for VFH and D38 for VFD which was similar to earlier published findings (Tanaka *et al.*, 2013, He *et al.*, 2010, Lu *et al.*, 2013b, Lu *et al.*, 2013a). This also indicated similar transport mechanism for both the transporters. Both the compounds ethidium bromide and norfloxacin fitted well in the central cavity of both the transporters. The only noticed difference was that while norfloxacin fitted in a straight fashion forming bonds with D40/38, Ethidium bromide showed bent fitting from 1^st^ to 7^th^ TM (Fig. 5).

### Ion binding and Electrostatic potential surface representation

MATE-type efflux pumps are secondary pumps that utilize Na^+^/H^+^ for transport (Jin *et al.*, 2014, Mohanty *et al.*, 2012). Earlier studies had proved that these were Na^+^ ion-driven transporters (Mohanty *et al.*, 2012). In the ion-dependence assays, we observed the transport capability was equally influenced by both sodium and potassium ions for VFH (reference). For better understanding of ion dependence and amino acid residues involved, we utilized IonCom server (https://zhanglab.ccmb.med.umich.edu/IonCom/) to predict their ion dependence and amino acid residues involved.

Results revealed that both the efflux pumps had no binding site detected for the following ions:(Cu, Fe, Ca, Mg, Mn, Na, K, CO, NO_2_, SO_4_, PO_4_). The predicted ion binding amino acid residues were D40, E51, Q82, D88, Y138, H163, N179, N309 and D393 for VFH and D38, H44, W49, Q81, N163 and D204 for VFD. Interestingly, docking studies clearly indicated that D38/40 of both the transporters were involved in binding. So, even though we couldn’t predict dependence of ion but D38/40 seems most plausible ion binding site for both the transporters which was similar to NorM from *V. cholerae* (He *et al.*, 2010).

Electrostatic potential of proteins is caused by charged amino acid residues in side chains, bound or unbound by various ions and play various roles such as protein folding and stability and function of the proteins. APBS pluggin in PyMol was used to calculate the electrostatic potential for both the proteins. It may reflects secondary and tertiary structures of transporters and their affinity to bind to substrates. Results revealed that both VFH and VFD had different hydrophobicity patches similar to other known MATE transporters (Fig 6a) (Zheng *et al.*, 2018). Both VFH and VFD formed a central cavity surrounded by charged electronegative amino acids. VFH also had an electropositive centre at the end of the cavity (Fig 6a). Subsequently, VFH and VFD with all possible interaction with norfloxacin via docking were mapped along and electrostatic potential was calculated, an interesting scenario came out. A sizable norfloxacin molecule was assembled in the cavity of VFH whereas fewer norfloxacin molecules were assembled in the cavity Of VFD. NorM which was taken as control had maximum norfloxacin molecules assembled in the cavity (Fig 6c). This was in sync with our earlier observation where VFH exhibited broader transport capability than VFD in MIC. The arrangement was mostly concentrated towards n-lobe which was similar like NorM (Lu *et al.*, 2013a)

**Fig. 6.**
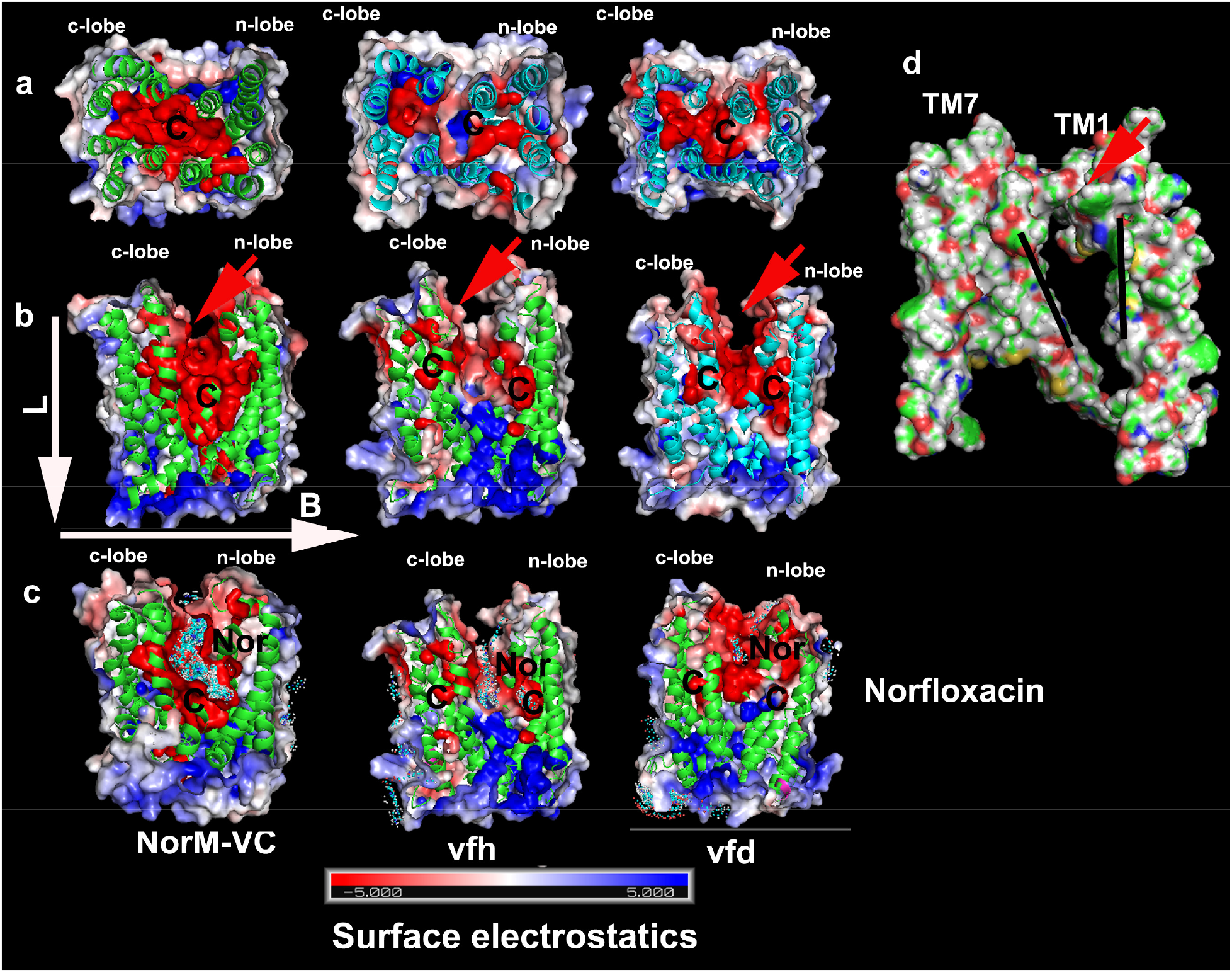
Surface charge differences between VFH, VFD and NorM-VC. Surface electrostatics were calculated by the APBS plugin in PyMOL. Cross-section model of VFH, VFD and NorM-VC surface representation is shown from the membrane region (a) or from the extracellular (b) sides (c) with norfloxacin and (d) Cross-section model of VFH/VFD exposing covalent interaction between TM1 and TM7. C represents cavity. Red arrows represent cavity.

It has been widely accepted that MATE transporters evolved from genetic duplication of flippases (Lomovskaya & Bostian, 2006, Tanaka *et al.*, 2013). Comparison of both the transporters with known structures of MATE transporters from *Camelina sativa*, NorM from *Vibrio cholarae* and MurJ flippases from *Thermosipho africanus* reveals interesting similarities and divergence from both VFH and VFD. D38/40 of both the transporters seemed to more conserved in bacterial transporters (Lu *et al.*, 2013b, He *et al.*, 2010) in comparison to their eukaryotic counterparts (Tanaka *et al.*, 2017). When both these transporters were compared to their evolutionary predecessors like MurJ (Kuk *et al.*, 2017), the difference is quite significant. MurJ flippases from *Thermosipho africanus* exhibited presence of 14 TMs arranged in 6+6 similar to bacterial transporters and functional role of last two TMs is unknown. Ser17, Arg18, Arg24, Arg52, and Arg255 in 1^st^ Tm of Murj has been proved important for recognizing the diphosphate and/or sugar moieties of lipid II whereas Asp in 1^st^ Tm of NorM has been proved for ion dependence and substrate transport (He *et al.*, 2010). It seems both VFH and VFD has similar topology like NorM with presence of D38/40 in 1^st^ TM creating a hydrophobic cavity for substrate recognition and transport.

During mapping of electrostatic potential of VFH and VFD, an aberrant architecture was also observed in both the transporters. MATE transporters usually have 12 TMs arranged in two bundles with 6 TMS each. Each forms one lobe and are usually free. The 1^st^ TM upon protonation displaces to 7^th^ TM to facilitate transport. But in case of both VFH/VFD, the loop connecting 1^st^ TM to 2^nd^ TM forms a covalent structure with loop of 7^th^ TM connecting 8^th^ TM as elucidated by surface model (Fig 6d). MATE type transporters are thought to be evolved of internal genetic duplication, resulting in two lobes. VFH/VFD had 10 TMs, still there was greater scope for transport. The formation of covalent bond reduces mobility of 1^st^ TM to be displaced to 6^th^ TM and may overall hinder transport efficiency (Fig 7b).

**Fig. 7.**
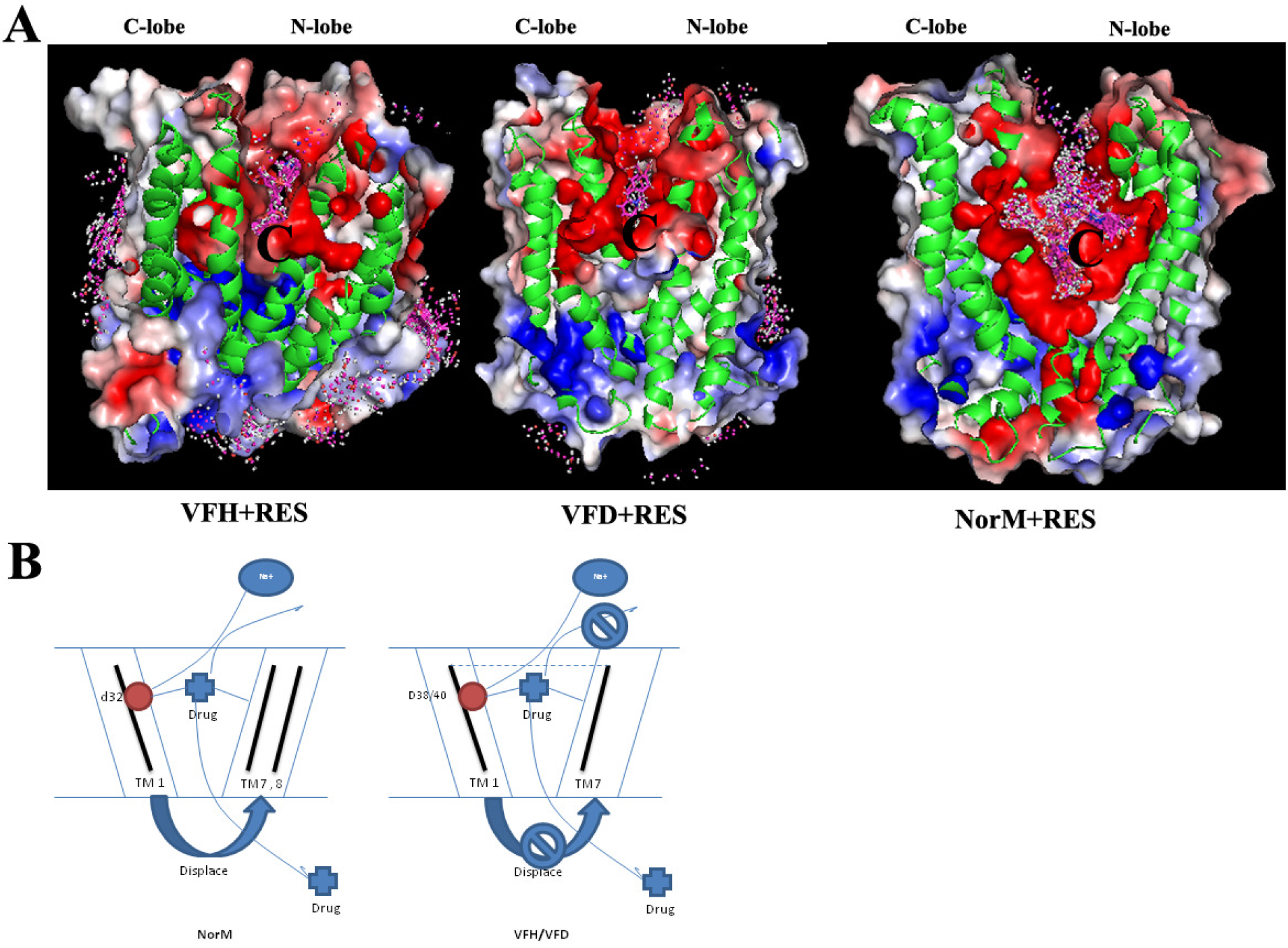
**A.** Docked pose of reserpine with *V. fluvialis* MATE efflux pumps (VFH and VFD). **B.** Proposed mechanism of transport of NorM-VC and VFD/VFH. Contrasting to NorM-VC, VFH/VFD forms covalent interactions between TM1 and TM7, which along low cavity volume leads to low transport capability of VFH/VFD.

### Mechanism of inhibition of both the transporters by reserpine

Our earlier observation with electrostatic mapping and inhibition of both the transporters with various EPIs had led to an indication that the transport of norfloxacin was indeed hampered in presence of these EPIs. Surprisingly, the similar effect couldn’t be observed in case of drug transport assay. Compounds like CCCP/FCCP, which are extremely toxic to cell at low concentration showed marginal inhibition. Reserpine which was fairly non-toxic, was able to inhibit both the transporters in real time drug transport assays. In our earlier study, we had observed 20 microgram/mL was sufficient to inhibit both the transporters (Mohanty *et al.*, 2012). Even in the current study, the MIC against the host was fairly high i.e >128 mcg/mL. So, range of concentration was quite enough to inhibit. Even thoughit has been earlier proved in case of Bmr transporters, where certain residues were implicated for inhibition (Klyachko *et al.*, 1997), no direct structural studies have ever proved the actual inhibition by reserpine in MATE efflux pump. To investigate the exact nature, we tried to dock reserpine with both the transporters along with NorM from *V. cholerae*.

Results revealed that reserpine could conveniently fit into electronegative cavities of both the transporters. VFH had 3 binding clusters with 19 binding sites whereas VFD had 2 binding clusters with 15 binding sites. Interestingly, NorM had 26 binding clusters with 190 binding sites.

In some of the earlier studies, MATE transporters have been co-crystalized along with the inhibitors. (Tanaka *et al.*, 2013). These studies have delineated the role of hydrogen bonding between transportersand ligand for exact inhibition mechanism. Our own observation with NorMhad hinted the same as there were almost 15 clusters and 96 binding sites forming hydrogen bonds in the cavity of NorM. But in case of VFH there was no predicted hydrogen bond formation and for VFD, 1 cluster and 3 binding site configuration was observed. Previous studies regarding co-crystallization of MATE-transporters with certain inhibitors had shed light on formation of electrostatic bonds for inhibition with verapamil and macrocyclic peptides (Radchenko *et al.*, 2015, Tanaka *et al.*, 2013). Observation from our earlier studies and current study reflect that reserpine acts pretty well at different concentrations also.

The exact mechanism of reserpine inhibition is binding to phenylalanine, tryptophan and tyrosine residues of Bmr efflux pumps (Steinfels *et al.*, 2004) and inhibition by reserpine does not involve ion bioenergetics of the cell and seems to be a non-specific inhibition of efflux pumps as was also seen in the current study. This led us to believe three important aspects of transport a. mobility of each lobe plays a vital role in transport b. size and electronegativity of central cavity plays important role for actual substrate recognition c. binding of ligand in electronegative pocket is not always electrostatic but may be covalent as reserpine was able to inhibit without actual hydrogen bond formation in both the transporters.

## Experimental procedures

### Cloning in pBAD-TOPO-TA vector

The primers used for cloning in pBAD vectors were VFH-BAD-F/VFH-BAD-R and VFD-BAD-F/VFD-BAD-R mentioned in supplementary table 2. The amplicons were subsequently purified using PCR purification kit (Qiagen) and ligated in pBAD vector (Invitrogen) according to manufacturer’s protocol.

### Expression and detection of recombinants harbouring pBAD efflux pump genes

pBAD clones for expression of VCH, VFH, VCD and *VFD* (where D- and H- are two MATE-type efflux pumps, VC stands for *V. cholerae* and VF stands for *V. fluvialis*) and Lac-His (positive control) were transformed in *E. coli* LMG194 and *E. coli* K-12 Δ *tolC* (JW5503-1, *E. coli* Genetic Resource Centre, Yale University) for expression as well as functional characterization. For Western blots, *E. coli* LMG194/K12 Δ*tolC* cells transformed with the recombinant vectors were grown in Luria-Bertani (LB) medium to an optical density of 0.5 at 600 nm (OD_600_) and induced with 0.2% L-arabinose for 4 hours and the total cell lysates prepared in SDS-PAGE lysis buffer were electrophoresed on 8% SDS-PAGE. Western blot was done with 0.45 micron nitrocellulose membrane (Whatman/Pall Life sciences) and blocked with 5% fat free milk powder in Tris buffered saline with 0.05% tween 20 (TBST) for 2 hours at room temperature. The blot was incubated with goat Anti-His/V_5_ primary antibody at 1:5000 dilutions in 0.5% milk powder in TBST for 2 hours at room temperature with constant shaking followed by washing in TBST and incubation with anti-Goat IgG-HRP (Horseradish Peroxidase)/Anti-Goat IgG-AP (Alkaline Phosphatase) (Jackson Immunoresearch) secondary antibody. The blots were developed either by using Novex ECL HRP chemiluminiscent development kit (Invitrogen) or by colorimetry using DAB/H_2_O_2_ (for HRP conjugated secondary antibody) or BCIP/NBT (Calbiochem) (for AP conjugated secondary antibody) according to the manufacturer’s instructions. For chemiluminiscence, the blot was dried and developed on X-ray.

### Detection of expression of recombinant proteins by confocal imaging

Briefly, the cells were grown and induced as discussed in earlier subsection of Experimental Procedures. 200 μL of induced cells were centrifuged at 5000 rpm for 5 min, the cells were resuspended in 50 μL of supernatant and were spotted on coverslips coated with 0.2% L-Poly-lysine (Sigma) in 24-well tissue culture plates (Corning) for 30 min at room temperature. Cells were then fixed using 4% paraformaldehyde (PFA) for 15 min followed by washing twice with 250 μL PBST (PBS containing 0.05% Tween 20) for 5 min without shaking. The cells were blocked with 5% Bovine Serum Albumin (BSA) in PBST for 1 hour at room temperature followed by washing thrice with 500 μL PBST for 10 min each and solutions were aspirated gently using a pipette. 50 μL mouse monoclonal anti-6X His tagged primary antibody (Calbiochem) was added at a dilution of 1:150 in 0.5 % BSA in PBST in the respective wells and incubated for 2 hours without shaking at room temperature. Washing was done thrice as described above. Goat anti-mouse secondary antibody labeled with Cy3 (Invitrogen) was added at the dilution of 1: 50 in 0.5 % BSA in PBST and incubated for 2 hours at room temperature followed by one washing step. Cells were counterstained with 200 μL of 4’, 6-diamidino-2-phenylindole (DAPI) at 1 μg mL^−1^ concentration and 50 μL of Film Tracer 1-43 membrane marker (Invitrogen) at 1 μg mL-1 concentration in DMSO. Confocal imaging was carried out using CTR 6500 confocal microscope (Leica). Diode lamp was utilised for detection of DAPI, UV HeNe lamp for CY3 and Argon lamp was used to detect membrane marker.

### Preparation of Membrane fraction

The membrane fraction was prepared according to the published protocol (Leyh, 1990). The *E. coli* LMG194/ K12 Δ*tolC* recombinant cells were grown and induced as described earlier. 25 mL of induced cultures were harvested at 5000 rpm, 4°C for 15 min. The supernatant was discarded and the pellet was washed with 5 mL ice cold PBS (phosphate-buffered saline) followed by centrifugation at 5000 rpm, 4°C for 10 min. The cells were resuspended in 1 mL lysis buffer (50 mM potassium phosphate, pH 7.8) having 1 mM phenylmethanesulfonyl fluoride and incubated in −80°C deep freezer for 2 hours. The samples were then thawed and sonicated 4 times with 30 sec on and off pulse on ice. The samples were then centrifuged at 8000 rpm, 4°C for 5 min. The supernatants were saved as soluble fraction and the pellet was treated with 200 μL of 2% sarcosyl solution (Sigma Aldrich) and incubated at room temperature for 30 min. The sample was then centrifuged at 15,000 rpm for 30 min at room temperature to obtain inner membrane proteins as supernatant and the rest of the proteins as pellet. Samples were subsequently subjected to Western blot for detection of recombinant proteins.

### Minimum Inhibitory Concentration (MIC) assays

The pBAD expression clones *of vfh*, *vch*, *vcd* and *vfd* were transformed in *E. coli* LMG194/K-12 Δ*tolC* cells, grown in LB broth containing ampicillin (100 μg mL^−1^) to an OD_600_ of 0.5 and induced with 0.2% L-arabinose for 4 hours. Kanamycin (30 μg mL^−1^) was additionally used for *E. coli* K-12 Δ*tolC* cells. Drugs/dyes were serially double diluted in LB broth containing 0.2% L-arabinose and 2 mL dilution/well were dispensed in 24 well tissue culture plates (Corning). OD_600_ of the induced sample was adjusted to 0.08-0.13 and 50 μL of this sample was added to each well. Incubation was done at 37°C. Readings were observed visually after 16 hours. The assays were repeated at least three times.

### Drug/Dye accumulation assays

For *E. coli* K-12ΔtolC cells harboring either pBAD-*vfh* or pBAD-*vfd* or not harbouring any plasmids were grown and induced as described earlier in Materials and methods. Ethidium bromide (1.9 μg mL^−1^) and norfloxacin (0.0156 μg mL^−1^) were used for accumulation assays. This was followed by single washing with 0.1 M Tris.HCl, pH 7.0 and resuspension in equal volume of the same buffer. The samples were collected every 15 min. The processing of the cells and monitoring of the fluorescent content was carried out exactly as described earlier (Mohanty *et al.*, 2012).

### Effect of Na^+^/K^+^ ions on efflux of drugs/dyes

*E. coli* K-12Δ*tolC* cells harboring either *pBAD-vfh* or *pBAD-vfd* or not harbouring any plasmids were grown in LB broth containing kanamycin (30 μg mL^−1^) and induced with 0.02% L-arabinose for 4 hours. Ampicillin was only added to the cells harbouring recombinant plasmids. The recombinants were grown and processed as described above in this section. The samples were then harvested by centrifugation, washed thrice with buffer containing 0.1M Tris.HCl, pH 7.0, and suspended in the same buffer to an OD_600_ of 1.0. The drug/dyes were used at 50% of the MIC values for transport studies. Norfloxacin at a final concentration of 0.0156 μg mL^−1^ (50% MIC concentration)along with 125 μg mL^−1^ of reserpine was added to the cell suspension and incubated for 30 min at 37°C. After 30 min, 1 mL samples were collected and stored in prechilled microcentrifuge tubes. Negative control (without drug/dyes) and EPI free control were also included in the study. Appropriate volume of 5 M and 0.5 M NaCl/KCl stock solution were added to assay mixture to a final concentration of 1M and 100 mM, incubated at 37°C for 15 minutes and 1 mL sample was collected. The volume was kept constant irrespective of salt concentration. Samples were then centrifuged at 10,000 rpm for 5 min at 4°C, washed once with 0.1 M Tris.HCl, pH 7.0 buffer and resuspended in 1 mL of 100 mM glycine.HCl, pH 3.0. These samples were subsequently processed and analyzed for fluorescence intensity as described in the previous sections. The maximum fluorescence observed in the experiment was considered to be 100%.

### Assays to determine efficacy of various EPIs on recombinant efflux pumps

The MIC of various EPIs was deduced against *E. coli* K-12 Δ*tolC*.Different concentrations of EPIs were prepared by two-fold dilution method in LB broth containing kanamycin (30 μg mL^−1^) and 100 μL/well was dispensed in 96-well tissue culture plates (Corning Inc.). The EPIs were procured from Sigma Aldrich. *E. coli* K-12 Δ*tolC* was grown in LB containing kanamycin (30 μg mL^−1^) to an OD_600_ of 0.08-0.13. Five microlitre of cells were inoculated in 100 μL medium containing various concentrations of EPIs in a 96 well plate. Observations were made visually after 16 hours. The experiments were repeated at least thrice. 50% concentration of MIC of each EPI was utilised in combination with various concentrations of norfloxacin. The *E. coli* K-12 Δ*tolC* carrying recombinant efflux pump plasmids and control *E. coli* K-12 Δ*tolC* were each grown in LB with kanamycin (30 μg mL^−1^) and ampicillin (100 μg mL^−1^). Overnight primary cultures were done in same way. Then 10 mL of secondary culture was prepared. Upon reaching OD_600_ of 0.5, the cells were induced with 0.02% L-arabinose for 4 hours with 200 rpm shaking at 37°C. The optical density of induced cells were adjusted to 0.08-0.13. Five microlitre of cells were inoculated in 100 μL medium containing 50% concentration of various EPIs along with various concentrations of norfloxacin in a 96 well plate (Corning Inc). Observations were made visually after 16 hours. The experiments were repeated at least thrice. The change in MIC was noted to observe efficacy of each EPI.

## Acknowledgments

The authors thank Indian Council of Medical Research, New Delhi for providing Senior Research Fellowship to PM (F/815/2011/ECD-II), Department of Biotechnology, New Delhi for Junior Research Fellowship (BT/PR/ 11634 INF/22/104/2008) of AS, Dr. A.K. Tiwari, Department of Developmental Biology, IIAR for helping in confocal imaging and the Puri Foundation for Education in India for support.

## Author contributions

Conceived and designed the experiments: AKB and PM. Performed the experiments: PM AS AKB. Analyzed the data: AKB PM. Contributed reagents/materials/analysis tools: AKB. Wrote the paper: AKB PM.

