## Supplementary Figure1, Table 1 and 2 for "Functional insights of two MATE transporters from *Vibrio fluvialis*": supplementary.docx

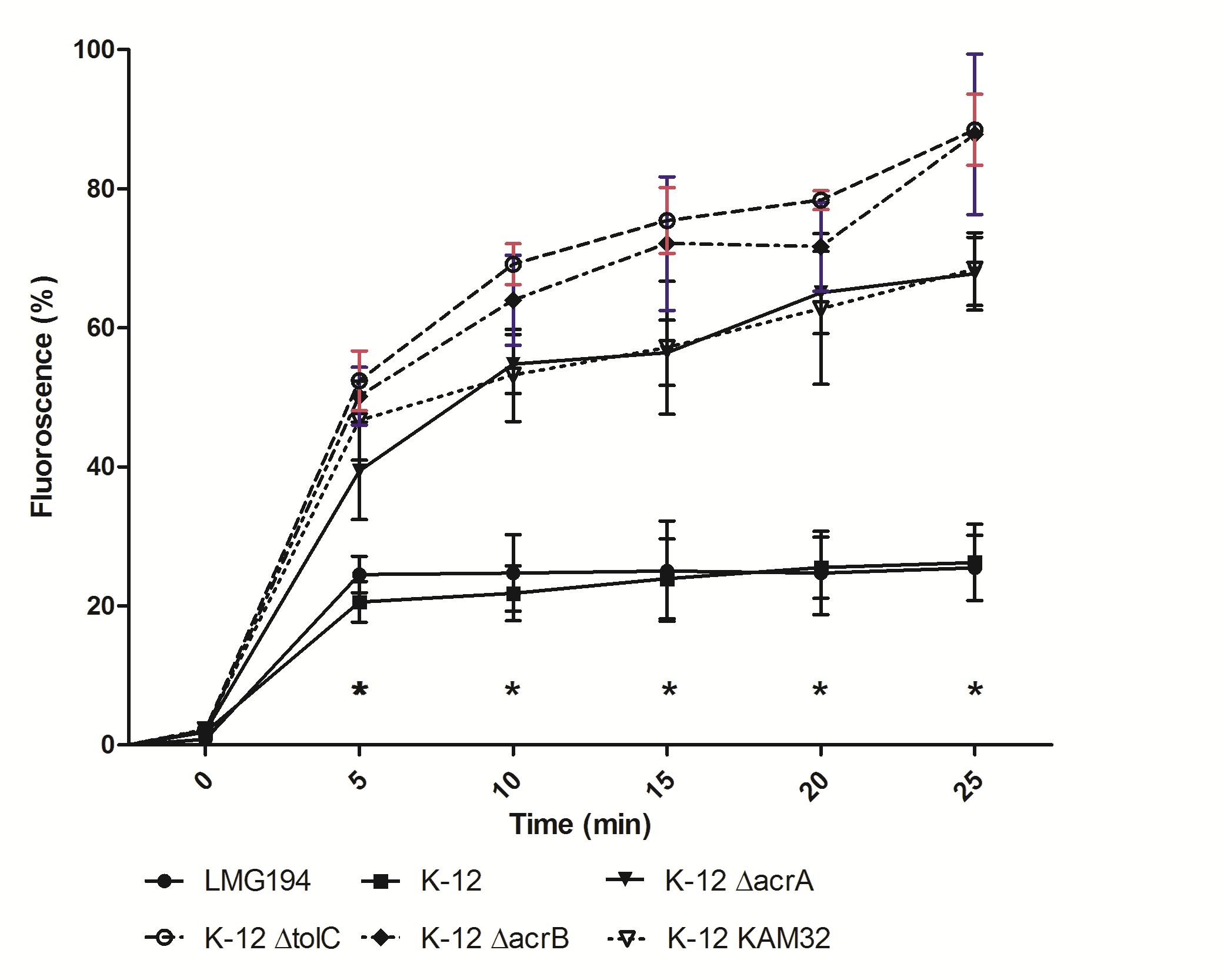

**Supplementary figure 1.** Levels of intracellular ethidium bromide concentration in various *E. coli* hosts. Accumulation was studied in the presence of 20 µg/mL ethidium bromide. Data is expressed as mean percentage fluorescence intensity of the maximum fluorescence intensity obtained during the experiment from three independent experiments and. *, the values of fluorescence for *E. coli* KAM32, *∆acrA*, *∆acrB* and *∆tolC* was significantly different as compared to LMG194 (*P* < 0.05).

**Supplementary Table 1. Optimisation of a hypersusceptible *E. coli* host for study of transport characteristics of recombinant efflux pumps**

|  | **MIC in µg /mL for the *E. coli* hosts** | | | | | |
| --- | --- | --- | --- | --- | --- | --- |
| **Drug/Strains** | **LMG194** | **K-12** | **KAM32** | ***∆acrA*** | ***∆acrB*** | ***∆tolC*** |
| Ethidium Bromide | 62.5 | 62.5 | 31.25-62.5 | 31.25 | 15.625-31.25 | <3.9 |
| **Trend** | ***∆tolC<∆acrB<∆acrA<* KAM32< LMG194/K-12** | | | | | |
| Norfloxacin | 0.25 | 0.25 | 0.125-0.25 | 0.0625 | 0.0625-0.125 | <0.0156 |
| **Trend** | ***∆tolC<∆acr<∆acrB<* KAM32< LMG194/K-12** | | | | | |
| Ciprofloxacin | 0.0125 | 0.00625 | 0.00625 | 0.003125 | 0.00625 | <0.00156 |
| **Trend** | ***∆tolC<∆acrA<∆acrB/*KAM32/ K12<LMG194** | | | | | |

*E. coli* K-12 (Wild type strain), LMG194 (genetically modified *E. coli* K-12 used for protein expression), *E. coli* KAM32 (strain with deletion of *acrB* and *ydhE*, which is used for functional characterisation of pBR322 recombinants), *∆acrA* (K-12 strain with deletion of connecting protein acrA of RND pump), *∆acrB* (K-12 strain with deletion of inner protein acrB of RND pump) and *∆tolC* (K-12 strain with deletion of outer membrane protein TolC of RND pump).

**Supplementary Table 2.** Primers used in the study

| **Primer Name** | **Sequences of primers (5’🡪3’)** |
| --- | --- |
| **VFH-BAD-F** | ATTTTCAATCAAATTCTTC |
| **VFH-BAD-R** | GTGTCTGTTGTCTCTCAAGTTTTATC |
| **VFD-BAD-F** | GTGTCACTGATTTTGCAAACGTT |
| **VFD-BAD-R** | AATGGAAAAGCGAGCCGG |
